# Formin FHOD1 regulates the size of EPEC pedestals

**DOI:** 10.1101/2020.06.15.149344

**Authors:** Xuyao Priscilla Liu, Mrinal Shah, Linda J. Kenney

**Author notes:** **Abbreviations:** Enteropathogenic *E. coli* (EPEC), Translocated intimin receptor (Tir), locus of enterocyte effacement (LEE), Attaching and effacing (A/E) lesions, Small molecule formin inhibitor (SMIFH2), Src family kinases (SFKs), wildtype (WT), standard error of the mean (S.E.M.).

## Abstract

Enteropathogenic *E. coli* (EPEC) is an extracellular pathogen that causes polymerization of actin filaments at the site of bacterial attachment, referred to as ‘actin pedestals’. Actin polymerization in the pedestal was believed to be solely regulated via the Nck-WASp-Arp2/3 pathway before formins were recently discovered to be associated with pedestals. Herein, we explored the collaborative role of formins in contributing to EPEC pedestal formation. In particular, we discovered that the formin FHOD1 preferentially localized to the pedestal base and its knockdown drastically reduced pedestal surface area. The pedestal localization of formin FHOD1 was found to be dependent on Tir phosphorylation at Y474, and on FHOD1 phosphorylation at Y99 from host Src family kinases (SFKs). Interestingly, differences in Arp2/3 and FHOD1 dynamics were observed. In large pedestals, Arp3 was nearly absent, but FHOD1 levels were high, suggesting that Arp2/3 and formins were segregated temporally. In line with this observation, as the pedestals grew in size, FHOD1 localization increased, while Arp3 localization decreased along the pedestals. Together, our results suggest that EPEC employs multiple actin nucleators that act at different stages of pedestal formation.

**Graphical abstract:** 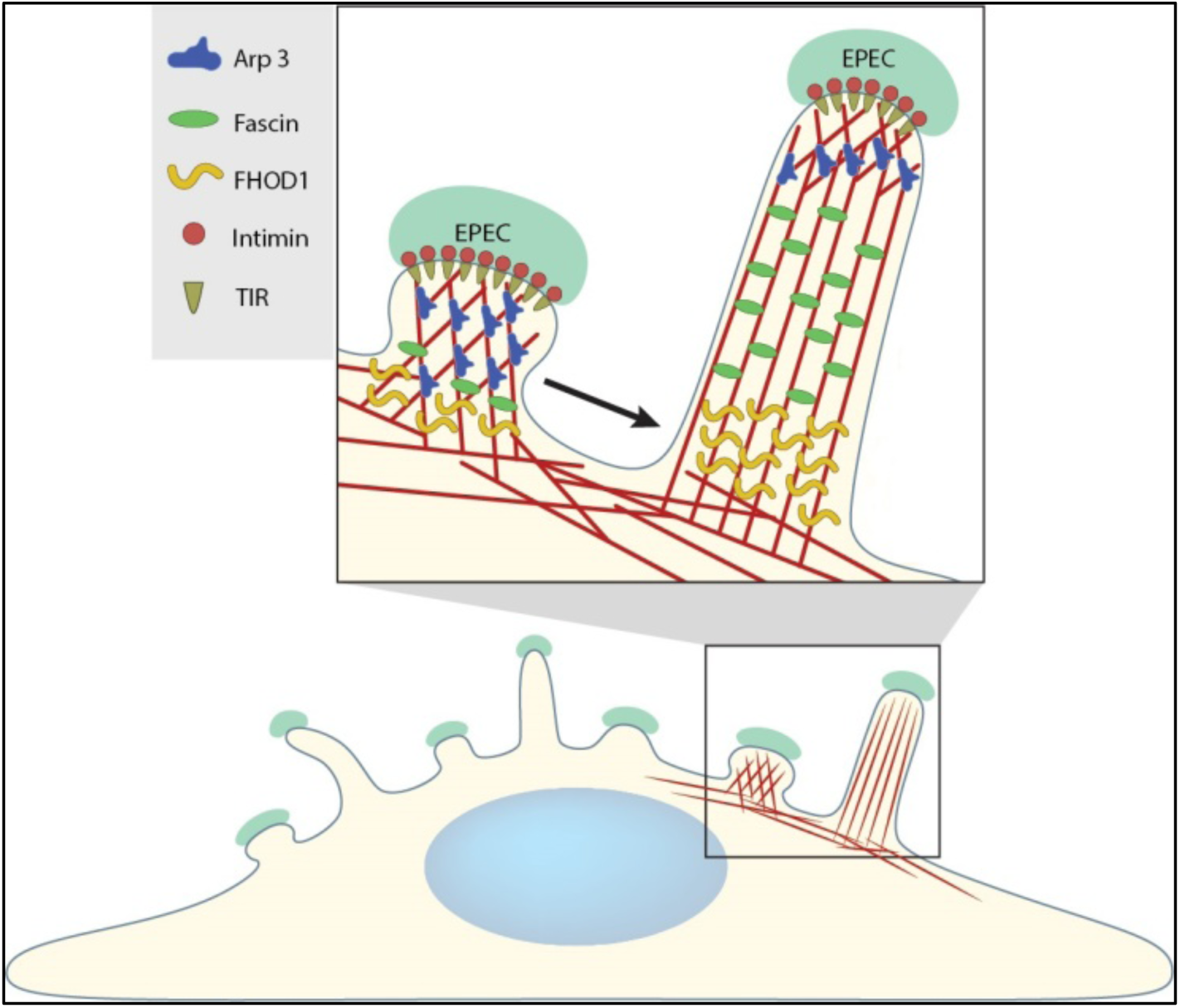

## Introduction

Enteropathogenic *E.coli* (EPEC) is a bacterial pathogen that evolved from *E. coli*, a commensal inhabitant of the gut, by the acquisition of pathogenicity islands and virulence genes (McDaniel, Jarvis, Donnenberg, & Kaper, 1995; McDaniel & Kaper, 1997; Tauschek, Strugnell, & Robins-Browne, 2002; Kaper, Nataro, & Mobley, 2004). EPEC is a leading cause of infantile diarrhea in developing countries, especially for children under 12 months of age (Nataro & Kaper, 1998; Kotloff et al., 2012; Kotloff et al., 2013; Croxen et al., 2013). The severity of the resulting illness is attributed to the diversity and complexity of the EPEC pathogen and its virulence factor repertoire, as highlighted in a recent comparative genomics study (Hazen et al., 2016). EPEC, and a closely-related pathogen Enterohemorrhagic *E. coli* (EHEC), have evolved sophisticated strategies to evade host immune responses, particularly phagocytosis, by intimately attaching to the extracellular surface of epithelial cells (Goosney, Celli, Kenny, & Finlay, 1999; Goosney, Gruenheid, & Finlay, 2000). Attachment leads to the formation of distinct ‘attaching and effacing’ (A/E) lesions on the gut epithelium characterized by microvillar effacement and dense actin-rich structures beneath the bacterial attachment site referred to as actin pedestals (Moon, Whipp, Argenzio, Levine, & Giannella, 1983; Taylor, Hart, Batt, McDougall, & McLean, 1986; Kaper et al., 2004).

The ability to form A/E lesions and actin pedestals is mainly encoded by the locus of enterocyte effacement (LEE). The LEE operon encodes a bacterial adhesin or intimin, several secreted proteins and a type III secretion system (T3SS), through which effector proteins are translocated into the host (McDaniel et al., 1995; Jarvis et al., 1995; Dean, Maresca, & Kenny, 2005; Dean & Kenny, 2009; Deng et al., 2012; Gaytán, Martínez-Santos, Soto, & González-Pedrajo, 2016). EPEC effectors alter host cell dynamics in multiple ways, with specific functions that are unique to each effector. Translocated intimin receptor, or Tir, is one of the first effectors to be translocated into the host cell (Kenny et al., 1997; Kenny, 1999; Mills, Baruch, Aviv, Nitzan, & Rosenshine, 2013). Tir is then inserted into the host plasma membrane where it then interacts with the bacterial surface protein intimin. This interaction acts as a trigger for the onset of EPEC pedestal formation (Kenny et al., 1997; Kenny & Finlay, 1997).

Using cultured epithelial cells for infection, it was discovered that tyrosine residue 474 was crucial for EPEC pedestal formation (Kenny, 1999). A redundant host kinase system, including Src-, Abl- and Tec-family kinases phosphorylates Tir (Phillips, Hayward, & Koronakis, 2004; Swimm et al., 2004; Bommarius et al., 2007). When Tir is phosphorylated at Y474, it interacts with the adaptor protein Nck1 and Nck2, which further interacts with N-WASP, bringing about Arp2/3-mediated actin polymerization (Kalman et al., 1999; Gruenheid et al., 2001; Lommel et al., 2001; Campellone et al., 2004). An additional tyrosine residue at Tir 454 was also shown to mediate weak actin polymerization independently of Nck (Campellone & Leong, 2005).

In addition to the well-established Tir:Arp2/3-complex actin pathway, the actin nucleator formin mDia1 was recently shown to form actin seeds and to promote Src family kinase activation for Tir phosphorylation in the formation of EPEC pedestals (Velle & Campellone, 2018). During their vast siRNA screen, knockdown of formin FHOD1 did not have a significant reduction in EPEC pedestal formation (Velle & Campellone, 2018). This result was unexpected, because formin FHOD1 was found to contribute in the actin-based motility of *Vaccinia virus* (Alvarez & Agaisse, 2013), and FHOD1 knockdown impaired *S.* Typhimurium invasion (Truong et al., 2013). For these reasons, we suspected formin FHOD1 was also involved in EPEC pedestal formation. In the current study, we examined the pedestal localization of formin FHOD1 during EPEC-induced actin rearrangement. Pedestal localization and effects on actin required phosphorylation of Src family kinases. In comparison to the Arp2/3 complex, formin FHOD1 was recruited to the pedestal at a much later time during infection. FHOD1 knockdown also reduced the pedestal length. Our results thus suggest that formin FHOD1 contributes to elongation of EPEC pedestals that are initiated from Arp2/3 complexes.

## Results

### Formins regulate EPEC pedestal formation in NIH3T3 cells

Although the general role of formins was reported to influence EPEC motility and colonization in NIH3T3 cells, a direct effect of formins on EPEC pedestals was not determined (Velle & Campellone, 2018). To fill that gap, we used the small molecule formin inhibitor SMIFH2 and pre-treated cells prior to infection with EPEC. Compared to untreated cells, we observed a drastic reduction in the surface area of pedestals formed in the presence of 25 µM SMIFH2 (Fig. 1A, 1C). The average area of pedestals formed in the presence of the inhibitor (16.20 ± 10.28 µm^2^) was reduced by nearly 50% compared to those formed in the absence of inhibitor (28.74 ± 14.99 µm^2^; see also Fig. S1). Furthermore, longer pedestals were almost completely absent; most pedestals formed in the presence of SMIFH2 were smaller than 20 µm^2^ (Fig. 1C). In contrast, the presence of the SMIFH2 had no significant effect on the number of pedestals formed compared to DMSO-treated control cells (Fig. 1B). Our results suggested two possibilities: that formins act as parallel actin nucleators in pedestals, thus affecting the kinetics of pedestal formation, or they could be involved in the pedestal elongation process, or both processes might be affected formins.

**Fig. 1.**
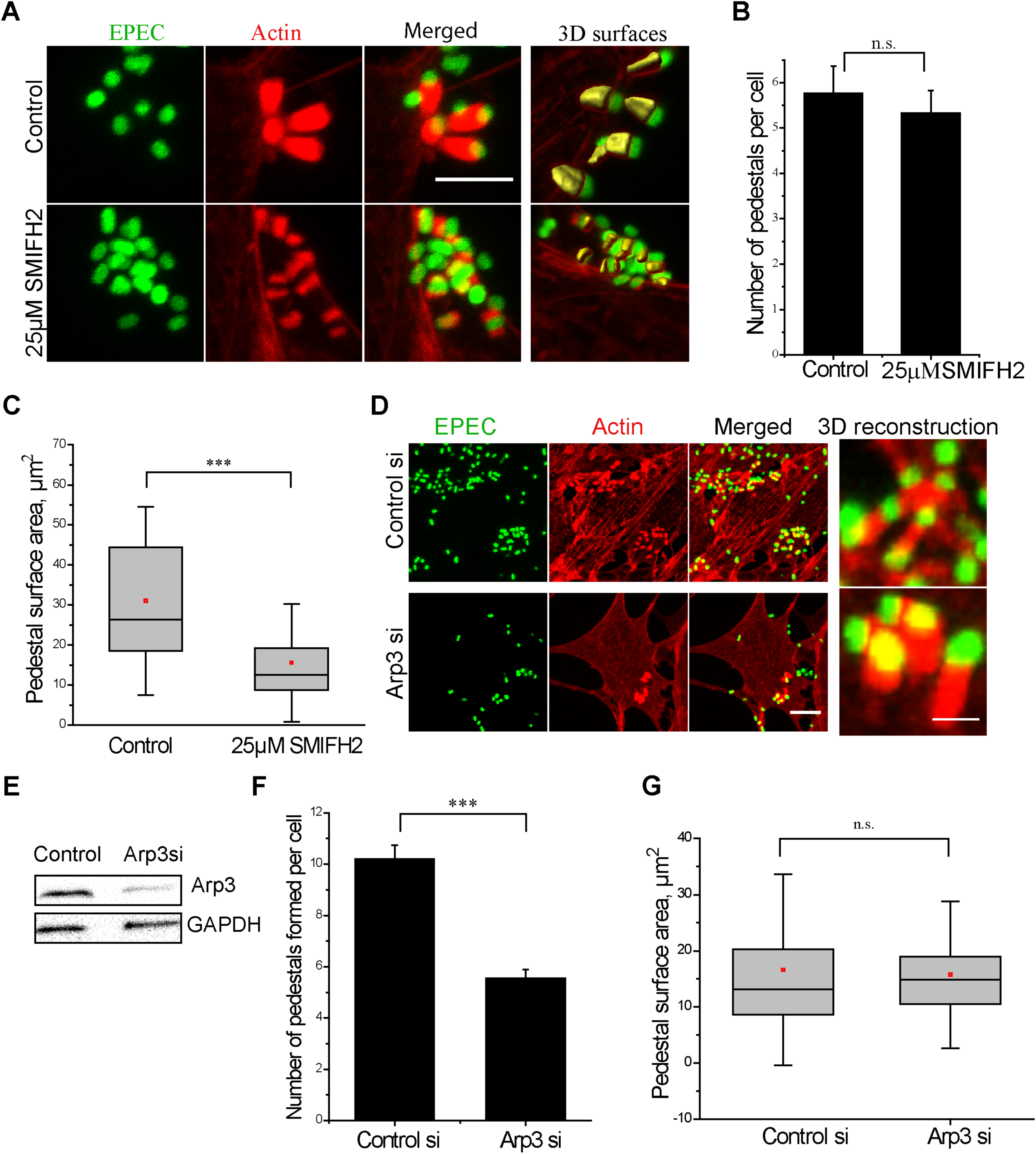
Formins regulate EPEC pedestal formation in NIH3T3 cells. **(A)** The top panel shows control cells treated with DMSO only. The left-most panel is EPEC-expressing GFP (green), next to it is Phalloidin 568-stained actin (red) and next to it the merged image is shown. Smaller pedestals were formed when NIH3T3 cells were pre-treated with 25 μM SMIFH2 (bottom panels). Scale bar = 5 μm. As shown in the right most panel, the surfaces of pedestals were marked using the actin channel (yellow) in the Imaris software and used for quantification. **(B)** The number of pedestals formed per cell did not change significantly upon SMIFH2 treatment. The mean ± S.E.M. is indicated. The graph represents the results of three independent experiments with in total 80-100 cells used for each group. **(C)** Quantification of the total surface area of the pedestals using Imaris, as shown in **(A)**. The total area of the pedestal was reduced significantly in the case of SMIFH2-treated cells. The graph represents the results of three, independent experiments with 100-150 pedestals used for each group. **(D)** Confocal Z-projected images of NIH3T3 cells infected with GFP-tagged EPEC (green). Actin was stained with Phalloidin-568. Arp3 knockdown led to formation of fewer pedestals (bottom panel) (see **(F)**). Scale bar = 10 μm. A zoomed region from the merged image reconstructed using Imaris, is shown in the right-most panel. Scale Bar = 2 μm. **(E)** A Western blot of Arp3 levels in NIH3T3 fibroblasts treated with a non-targeting siRNA (Control) or Arp3siRNA; GAPDH was used as a loading control. **(F)** The number of pedestals formed per cell decreased upon Arp3 knockdown at 5 hours post EPEC infection. Note that the number of pedestals formed was only reduced by 50%. The mean ± S.E.M. is indicated. The graph represents three, independent experiments with in total 80-100 cells used for each group. **(G)** Arp3 knockdown did not affect the total pedestal area, which remained unchanged compared to the control. In the box plots, the values of the median (black line), the mean (red dot), lower and upper quartiles (box) and the standard deviation (whiskers) are indicated. The significance of the difference between groups was estimated by two-tailed Student’s *t* test. n.s., P > 0.05 (not significant); **, P ≤ 0.01; ***, P ≤ 0.001; ****, P ≤ 0.0001.

Arp2/3 was previously identified as the major actin nucleator in EPEC pedestals (Kalman et al., 1999), but CK666 treatment (50 µM) only partially reduced the formation of the pedestals (Fig. S2). Transcriptional silencing of Arp3 reduced the number of pedestals formed by only ∼54% (Fig. 1D-F), which was similar to previous observations (Velle & Campellone, 2018). Thus, a significant population of pedestals was still visible when Arp2/3 was substantially reduced. Interestingly, the pedestals that were formed were similar in surface area to the wild type pedestals (Fig. 1G). This result suggested that in addition to the canonical Nck-Arp2/3 pathway, an alternate actin polymerization pathway involving formins drives EPEC pedestal formation in NIH3T3 cells.

### Formin FHOD1 localized to the base of EPEC pedestals

Because we observed a reduction in the length of EPEC pedestals using a general formin inhibitor (Fig. 1), we wanted to identify which specific formin was strongly localized to the pedestal sites. GFP-tagged constructs of formin proteins, including FHOD1, DAAM1, FMNL3, INF2, mDia1 and mDia2, were transfected separately into NIH3T3 cells, which were then infected with EPEC. The fluorescence and immune-stained images indicate that pedestals were formed beneath EPEC in all cases, but the distribution of each formin protein was distinct, and the intensity level varied as well (Fig. 2). GFP-tagged FHOD1 was recruited to actin-rich pedestal sites. It was distributed along the length of pedestals but was concentrated at/near the pedestal base (Fig. 2A). GFP-tagged formin DAAM1, FMNL3 or INF2 were not recruited to pedestal sites, instead they showed a homogeneous, cytoplasmic distribution (Fig. 2B, top 3 rows). GFP-tagged formin mDia1 and mDia2 were recruited to EPEC pedestal sites (Velle & Campellone, 2018), but compared to FHOD1, the fluorescence signals of mDia1 and mDia2 were relatively diffusive and were not concentrated predominantly at the base of EPEC pedestals (Fig. 2B, bottom 2 rows). The diffusive pedestal localization of mDia1 in NIH3T3 cells that we observed differed from previous reports where a localized accumulation of mDia1 at the pedestal was reported in HeLa cells (Velle & Campellone, 2018). Thus, among the six formin proteins we tested, formin FHOD1 localized strongly to the base of EPEC pedestals, and mDia1 & mDia2 showed a slight recruitment to pedestals, whereas the formins DAAM1, FMNL3, INF2 were not recruited to EPEC pedestals (Fig. 2B). Moreover, immune-staining of endogenous FHOD1 with a specific antibody also localized FHOD1 to EPEC pedestals (Fig. S2A). Thus, in NIH3T3 cells, formin FHOD1 is the major formin that is strongly localized to EPEC pedestals.

**Fig. 2.**
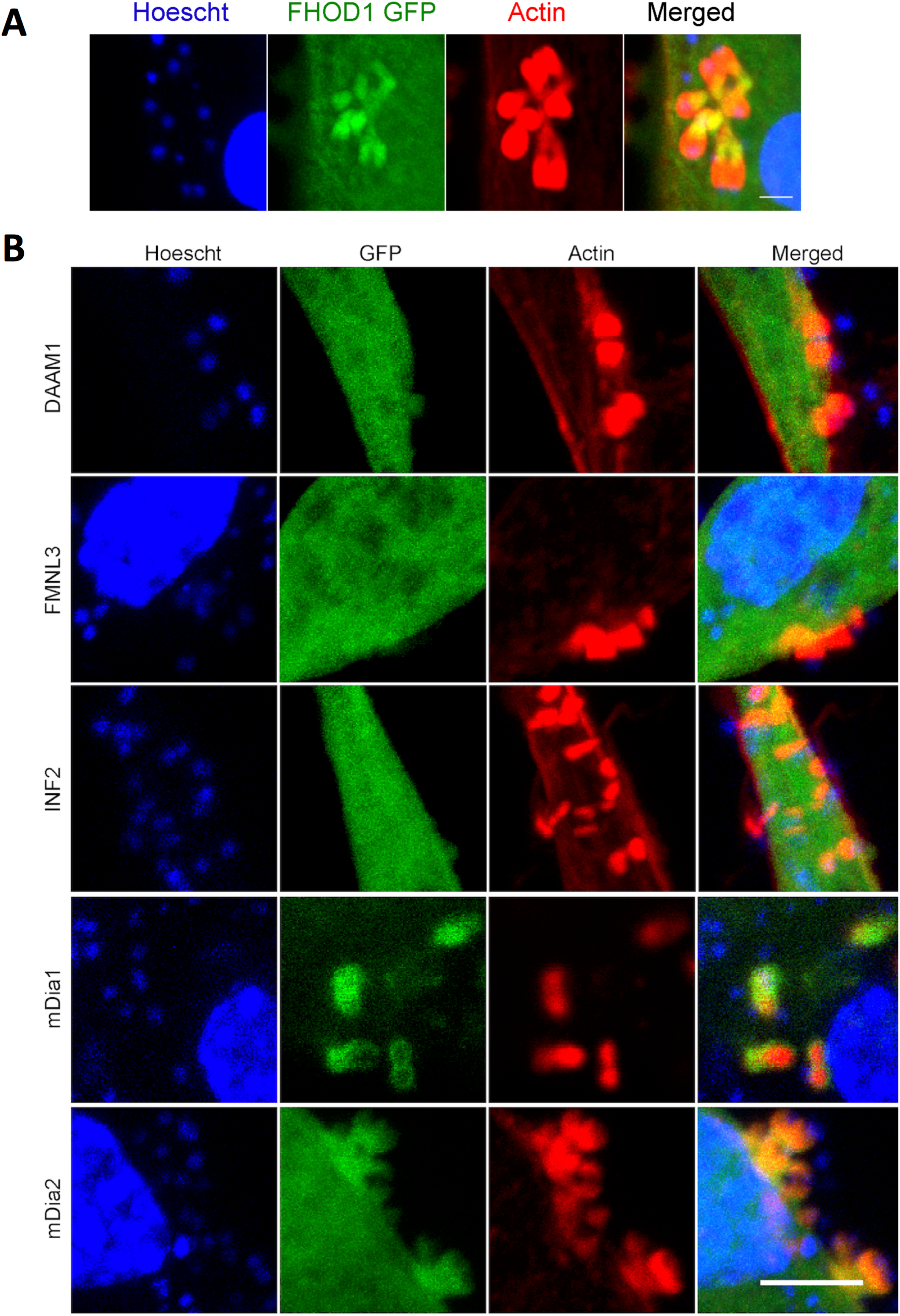
Formin FHOD1, and not formins DAAM1, FMNL3 and INF1 was localized to the base of EPEC pedestals. **(A)-(B)** Each panel shows NIH3T3 cells transfected with GFP-tagged constructs of **(A)** FHOD1, **(B)** DAAM1, FMNL3, INF2, mDia1 and mDia2, respectively and infected with EPEC. As shown in **(A)**, FHOD1 shows a strong localization to the base of the pedestals. Scale bar = 2 μm. As shown in the bottom two panels of **(B)**, mDia1 and mDia2 showed slight localization to pedestals (compare GFP and actin), whereas DAAM1, FMNL3 and INF1 (top three panels) did not localize to the pedestals. Actin was stained with Phalloidin-568. Bacteria and host cell nuclei were stained with Hoechst. Scale Bar = 5 µm.

### Formin FHOD1 promotes EPEC pedestal elongation

We next used RNAi silencing to examine how the absence of FHOD1 specifically affected pedestal formation. We used a pool of four independent siRNAs targeted to four different sites in the FHOD1 gene to achieve silencing. The numbers of pedestals formed were not significantly reduced in the absence of FHOD1 (Fig. 2A, 3A), although pedestals were significantly reduced in size (Fig. 2A, 3B). In addition to the siRNA pool, we also examined the effect of each siRNA individually. Knockdown of FHOD1 using individual siRNAs led to a reduction in pedestal size, but the effect varied depending on the amount of FHOD1 remaining (Fig. S3A-C). This result was consistent with our results using the formin inhibitor SMIFH2 (Fig. 1). Interestingly, a double knockdown of both FHOD1 and Arp3 resulted in an additive phenotype of both smaller and fewer pedestals (Fig. 3A-C).

**Fig. 3.**
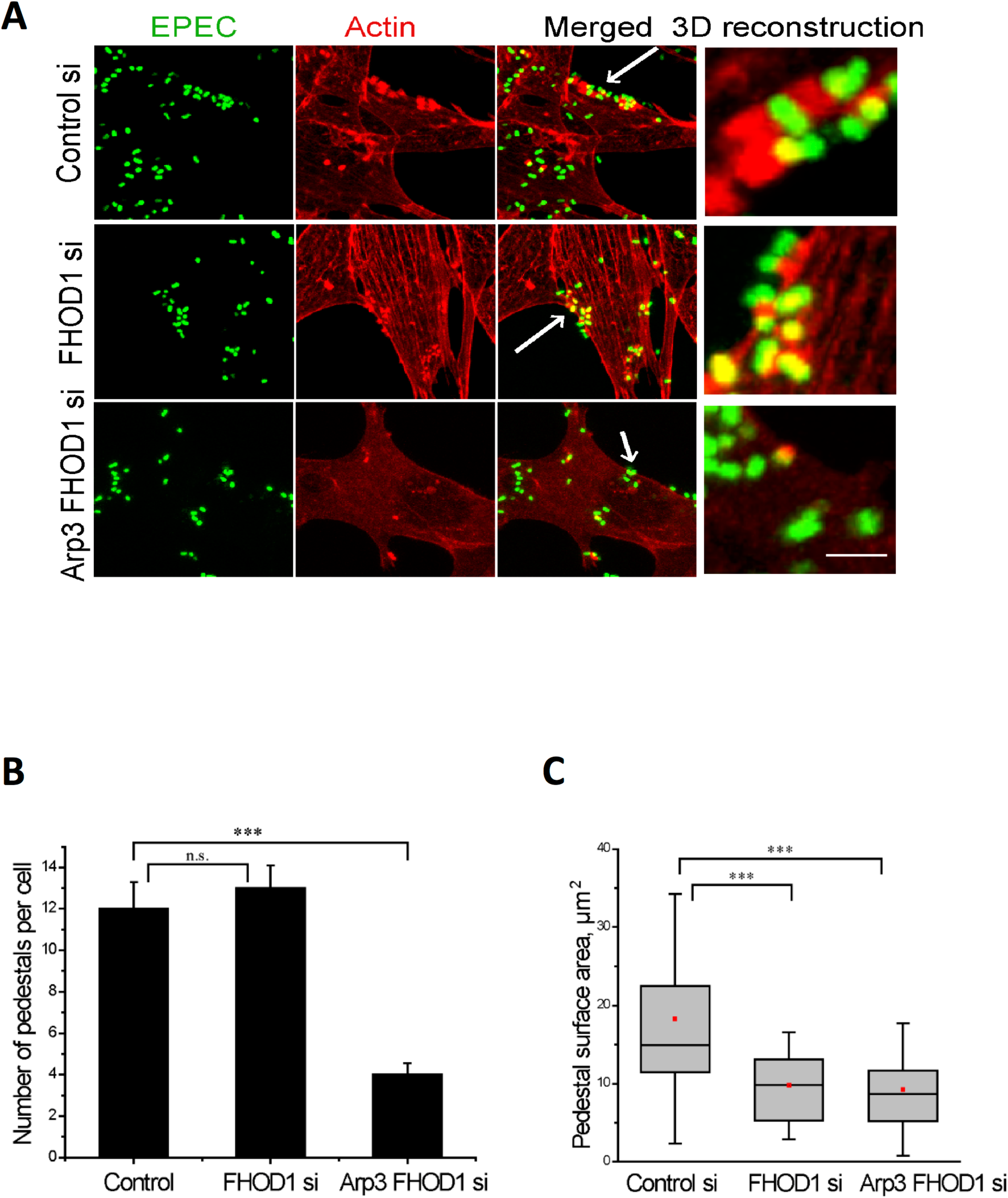
Formin FHOD1 controls EPEC pedestal size. **(A)** Smaller pedestals were formed when FHOD1 was knocked down (middle panels), compared to a non-targeting siRNA treated control (top panels). Smaller and fewer pedestals were formed when both Arp3 and FHOD1 were knocked down (bottom panels). Actin was stained with Phalloidin-568 (red); EPEC was GFP-tagged (green). The arrows point to the pedestals formed at the sites of GFP-tagged bacteria. Scale bar = 10 μm. The right-most panel shows a zoomed area from the merged image, reconstructed using Imaris. Scale Bar = 2 μm. **(B)** The number of pedestals formed per cell remained unaffected during FHOD1 knockdown, whereas they were significantly decreased in the Arp3/FHOD1 double knockdown. The graph represents the results of two, independent experiments with in total 80-100 cells used for each group. The mean ± S.E.M. is indicated. **(C)** Quantification of the total surface area of the pedestals using Imaris showed that the average pedestal area was reduced significantly (50%) when FHOD1 was reduced and was similar to the level in the Arp3 FHOD1 double knockdown. The graph represents results from two, independent experiments with 100-150 pedestals used for each group. In the box plots, values of the median (black line), the mean (red dot), lower and upper quartiles (box) and the standard deviation (whiskers) are indicated. The significance of the difference between groups was estimated by a two-tailed Student’s *t* test. n.s., P > 0.05 (nonsignificant); **, P ≤ 0.01; ***, P ≤ 0.001; ****, P ≤ 0.0001.

### Phosphorylation of FHOD1 is essential for its localization

Formin FHOD1 is recruited to focal adhesions by phosphorylation of a C-terminal tyrosine residue, and then activated by the Rho-associated protein kinase (ROCK) (Gasteier et al., 2003; Iskratsch et al., 2013). Thus, we suspected that phosphorylation might be important in activation or localization of formin FHOD1 at EPEC pedestals. We transfected the GFP-tagged construct of FHOD1 containing a tyrosine substitution (FHOD1 Y99F) into NIH3T3 cells and compared its localization with the wild type FHOD1 construct (Fig. 4A-B). Localization was quantified by comparing the ratio of intensities of FHOD1 at the pedestal to the average FHOD1 intensity in the cell. The intensity of wild type FHOD1 increased ∼2-fold at the pedestals, whereas the intensity of Y99F-FHOD1 was not localized to the pedestals, i.e. the intensity of GFP fluorescence was the same as an untagged GFP (Fig. 4B). This result suggested that tyrosine phosphorylation was required for FHOD1 localization to the pedestals.

**Fig. 4.**
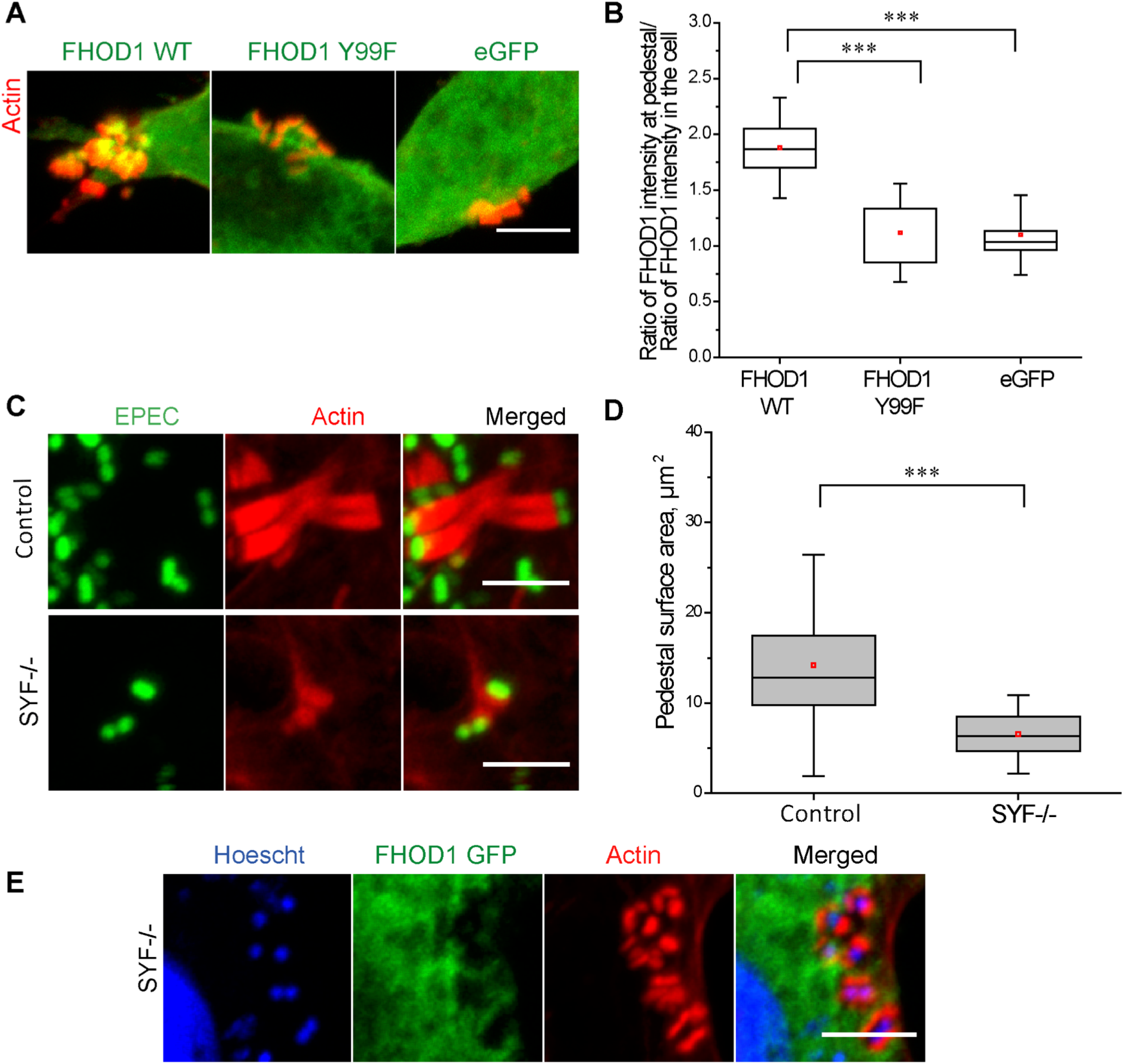
Phosphorylation of FHOD1 is essential for its localization. **(A)** FHOD1 Y99F (green) did not localize to the pedestals, and was similar to the negative control eGFP. Actin was stained by Phalloidin-568 (red). **(B)** Quantification of the ratio of intensity of FHOD1 at the pedestal versus the intensity in the cell indicated that the FHOD1 Y99F mutant did not localize to the pedestals as compared to WT FHOD1. The graph represents the results of two, independent experiments with 100-150 pedestals used for each group. **(C)** Smaller pedestals were formed in a SYF (-/-) fibroblast cell line compared to SYF (+/+) cells. Actin (red) was stained with Phalloidin-568 and EPEC-GFP (green) was used for infection. Scale Bar = 5 μm. **(D)** Quantification of the total area of the pedestal using Imaris showed that it was reduced by 55% in SYF (-/-) fibroblasts compared to SYF (+/+). The graph represents the results of three, independent experiments with 100-150 pedestals used for each group. In the box plots, the values of the median (black line), the mean (red dot), lower and upper quartiles (box) and the standard deviation (whiskers) are indicated. The significance of the difference between groups was estimated by two-tailed Student’s *t* test. n.s., P > 0.05 (not significant); **, P ≤ 0.01; ***, P ≤ 0.001; ****, P ≤ 0.0001. **(E)** This panel shows WT FHOD1-GFP transfected in SYF (-/-) fibroblasts (Iskratch *et al.*, 2013). The WT FHOD1-GFP (green) did not localize to the pedestals in SYF (-/-) cells. Actin (red) was stained with Phalloidin-568 and host nuclei and bacteria were stained with Hoechst. Scale Bar = 5 μm. Mean ± S.E.M. are indicated.

The EPEC effector Tir is phosphorylated on tyrosine residues by several redundant host kinases, including: Src family kinases Abl and Arg (Swimm et al., 2004). We therefore investigated whether these same kinases might be involved in the activation of FHOD1. We infected mouse fibroblasts deficient in Src, Yes, and Fyn (SYF-/- cells) (Klinghoffer, Sachsenmaier, Cooper, & Soriano, 1999; Phillips et al., 2004) and compared them to SYF-/- cells that were stably transfected with Src, Yes and Fyn. EPEC infection of SYF-/- cells resulted in significantly smaller pedestals compared to the control cells (Fig. 4C-D). We then transfected wild-type FHOD1 into SYF-/- cells, to determine whether SYF kinases were involved in its activation. Wild type FHOD1 no longer localized to the pedestals in SYF-/- cells (Fig. 4E). Taken together, these results indicate that phosphorylation was essential for the localization of FHOD1 to pedestals, and phosphorylation was mediated by redundant Src family kinases.

### Fyn tyrosine kinase recruits formin FHOD1 to EPEC pedestals

Since SYF-/- cells were unable to recruit FHOD1 to EPEC pedestals (Fig. 4E), we re-introduced Src or Fyn individually into SYF-/- cells and infected them with EPEC. The localization of FHOD1 to the pedestals was then examined (Fig. 5). When SYF-/- cells over-expressing Src-mCherry were infected with EPEC, actin-rich pedestals were formed under the pathogen as usual, but FHOD1 did not accumulate at the pedestal sites (Fig. 5A, middle panel). In contrast, when SYF-/- cells over-expressing Fyn-mCherry were infected with EPEC, FHOD1-GFP accumulated near the base of actin-rich EPEC pedestals (Fig. 5A, bottom panel), as observed previously (Fig. 4). The expression of Src or Fyn was confirmed (Fig. 5A, right panels). Fyn-mCherry co-localized with actin pedestals, whereas Src-mCherry did not.

**Fig. 5.**
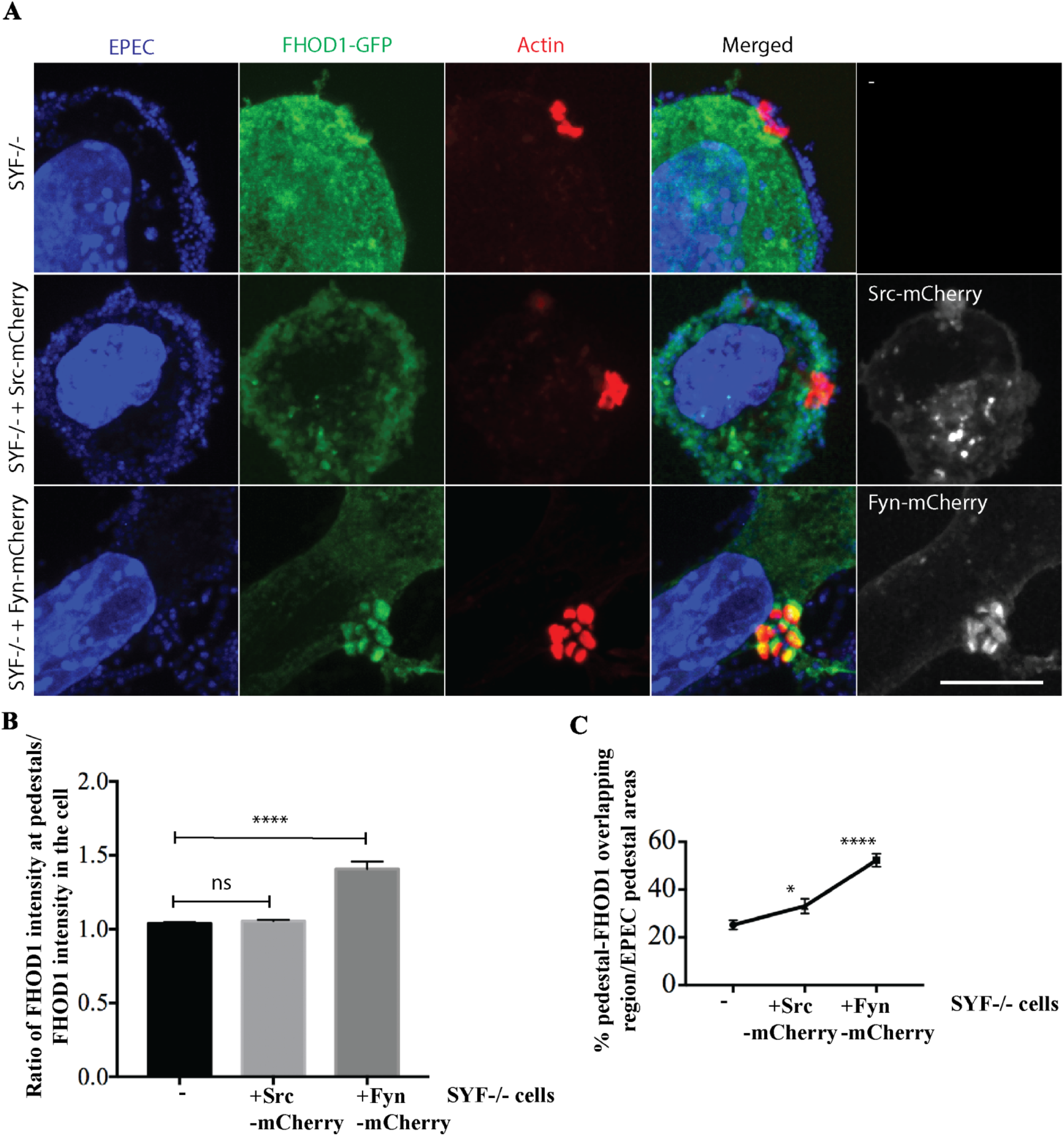
Src family kinase Fyn accounts for the recruitment of formin FHOD1 to EPEC pedestal. **(A)** The confocal images (Z projection, Sum intensity in Fiji software) of SYF-/- cells transfected with FHOD1-GFP and infected with EPEC for 5 hours of infection. The staining of nucleus and bacteria by DAPI is shown on the left column, followed by FHOD1-GFP, and actin filaments stained by phalloidin 680 and the merged images are shown to the second right cloumn. The overexpression of Src-mCherry or Fyn-mCherry in SYF-/- cells was verified and are shown in the last column to the right. FHOD1 colocalizes with actin-rich pedestals in Fyn-mCherry rescued SYF-/- cells. **(B)** The ratio of FHOD1-GFP intensity at pedestals and inside the cells is plotted. Higher degree of FHOD1-GFP accumulation was found to the pedestals in Fyn-added back cells. **(C)** The percentage plot of FHOD1-actins overlapping areas over EPEC pedestals is shown to show the recruitment of FHOD1-GFP to the pedestals. See **Materials and Methods**. The mean ± S.E.M. is indicated. Three independent experiments were included for statistics. Scale bar = 10 μm. ns, not significant; *, p<0.1; ****, p<0.0001.

The pedestal localization of FHOD1-GFP was then quantified (see Methods), the results are shown in Fig. 5B-C. In the absence of Src family kinases, the average FHOD1-GFP intensity in pedestals was equivalent to the average FHOD1-GFP intensity in the cytoplasm (ratio =1). This ratio remained at 1 when Src was re-introduced into SYF-/- cells. However, when Fyn was re-introduced to SYF-/- cells, the ratio increased by 40% (to 1.4, Fig. 5B). To corroborate this result, we compared the intensity of FHOD1-GFP and actin in the pedestals of SYF-/- cells in which Fyn had been reintroduced (see Methods). There was a ∼2-fold increase in intensity compared to SYF-/- cells, as shown in Fig. 5C. When Src was reintroduced into SYF-/- cells, the intensity only increased ∼1.3-fold (Fig. 5C). Thus, Fyn plays the major role in recruiting FHOD1 to the pedestals, although Src appears to make a minor contribution (see Discussion).

### FHOD1 localization to the pedestals depends on Tir tyrosine residue 474

Previous studies reported that Arp3 was localized underneath EPEC through an interaction with Tir in a process that required phosphorylation (Gruenheid et al., 2001). We wanted to determine whether phosphorylation of Tir was essential for FHOD1 localization, as reported for Arp3. Tir has four tyrosine residues that can be phosphorylated, two of them (Y454 and Y474) are directly linked to actin polymerization (Gruenheid et al., 2001; Campellone et al., 2004; Campellone & Leong, 2005). We used EPEC Tir tyrosine mutant strains: EPEC Y474A, EPEC Y454A and EPEC Y454AY474A (a gift from Gad Frankel; Crepin et al., 2010). Upon infection with WT EPEC, FHOD1-GFP was recruited to the pedestals (Fig. 6A, 1^st^ row). However, when NIH3T3 cells were infected with EPEC Y474A, FHOD1-GFP did not accumulate at the pedestal base (Fig. 6A, 2^nd^ row). NIH3T3 cells infected with EPEC Y454A were similar to the WT (Fig. 6A, 3^rd^ row). Infection with the double mutant Y454A-Y474A, showed a similar diffusive distribution of FHOD1-GFP as the single Y474A mutant (Fig. 6A, last row). Enlarged figures are shown in Fig. 6B for each condition. The ratio of FHOD1-GFP intensity at pedestal sites and inside cells was plotted in Fig. 6C. An averaged ratio of FHOD1 intensity of 1.2 was determined for both EPEC WT and EPEC Y454A, compared to FHOD1-GFP intensity in the cytoplasm. In the absence of Tir Y474, this ratio decreased to 1 (Fig. 6C). Thus, EPEC Tir Y474 is required for concentrating FHOD1-GFP at the pedestals, as reported for Arp2/3.

**Fig. 6.**
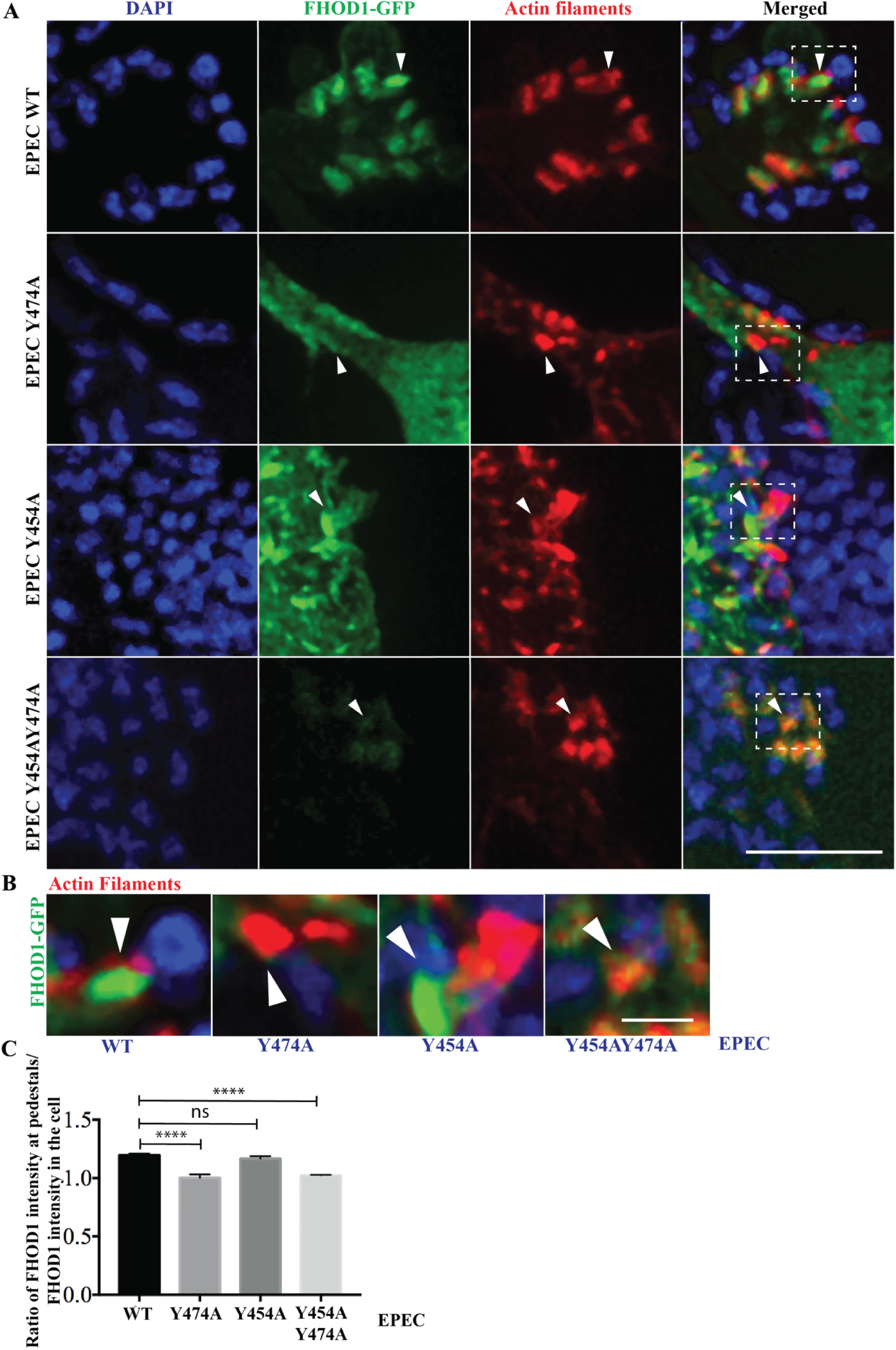
EPEC Tir Y474 promotes the recruitment of formin FHOD1 to EPEC pedestals. **(A)** The super-resolution SIM images (Z projection, SUM intensity in Fiji software) of NIH3T3 cells transfected with FHOD1-GFP and infected with various EPEC strains for 5 hours of infection. The staining of nucleus and bacteria by DAPI is shown on the left, followed by FHOD1-GFP, and actin filaments stained by phalloidin 680, the merged images are shown to the right. A white dotted box is drawn on the merged images with white arrowheads highlighting the representative EPEC pedestals induced by various EPEC strains. Those white boxes are enlarged in (**B)** to show the recruitment of FHOD1 to the pedestal sites. Scale bar = 5 μm. **(B)** Enlarged merged images from **A** with white arrowheads pointing to the representative actin aggregated induced beneath EPEC strains. FHOD1-GFP was accumulated at the pedestals induced by EPEC WT and EPEC Y454A strains, unlike in the cases of EPEC Y474A and EPEC Y454AY474A. Scale bar = 1 μm. **(C)** The ratio of FHOD1-GFP intensity at pedestals and inside the cells is plotted. See **Materials and Methods**. Three independent experiments were included for statistics. The mean ± S.E.M. is indicated. ns, not significant; ****, p<0.0001. EPEC strains are in blue, FHOD1-GFP in green and actin filaments in red. SIM: structured illumination microscopy.

### Arp3 and FHOD1 localization to pedestals is heterogeneous

Our results demonstrated a strong localization of formin FHOD1 at the base of the pedestals (Fig. 2). We next examined Arp2/3 complex localization with respect to FHOD1 to determine whether their localization was spatially distinct. Co-transfection of Arp3-mCherry with FHOD1-GFP identified an intense band of Arp3 localized near the bacterial end of the pedestal (Fig. 7A), whereas a dense patch of FHOD1 was observed at the opposite end, i.e., at the pedestal base. The localization of endogenous Arp3 was also observed by staining with Arp3 antibody (Fig. 7B). Surprisingly, pedestals were heterogeneous in size and they exhibited significant differences in the amount of Arp3 localized (Fig. S5). Overall, a higher intensity of Arp3 was associated with newly formed pedestals and a lower intensity was associated with older pedestals.

**Fig. 7.**
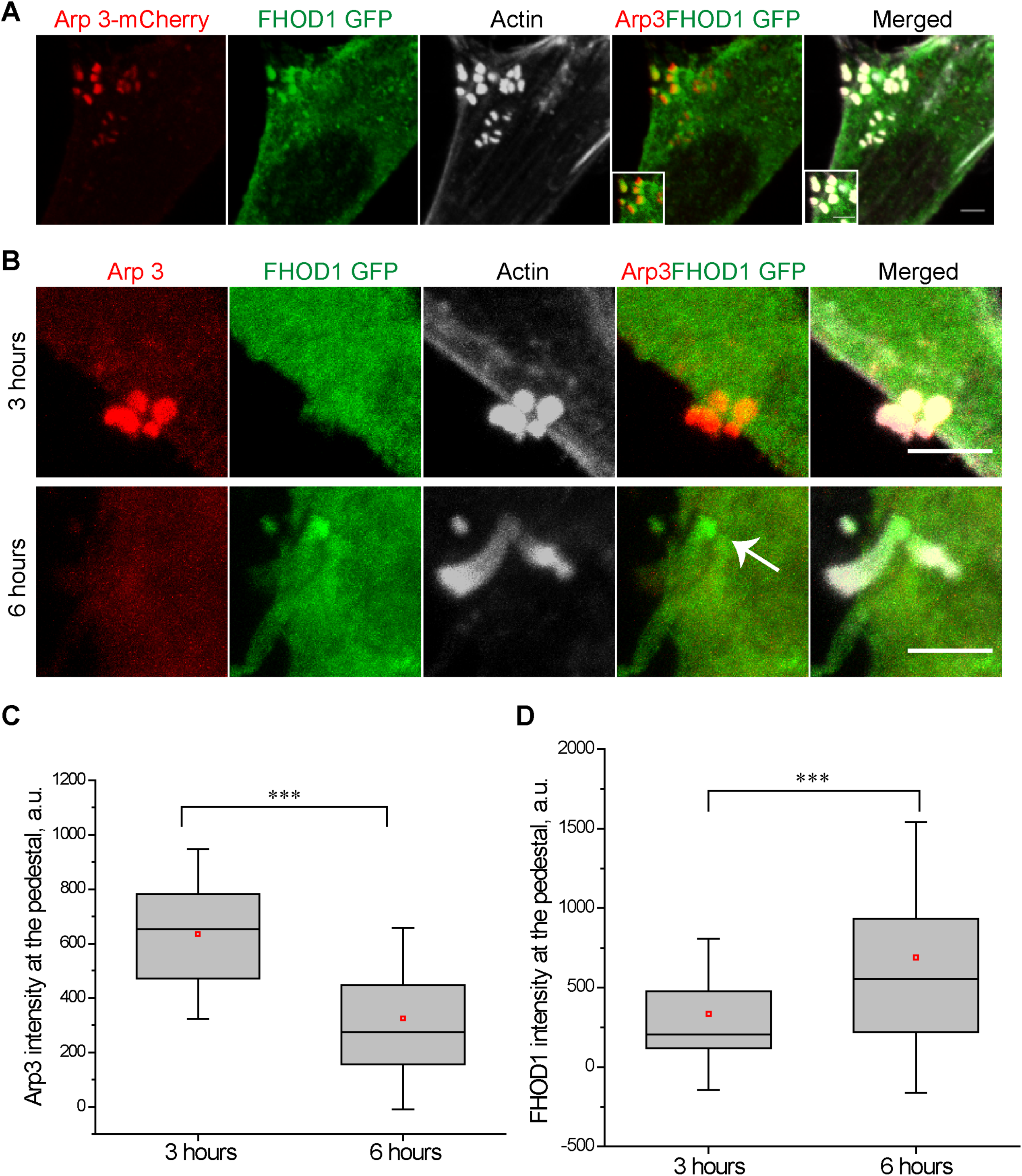
Arp3 and FHOD1 localization to the pedestals was heterogeneous. **(A)** The panel shows the localization of Arp3-mCherry and FHOD1-GFP to the pedestals. Arp3 (red) localized near the bacterial side of the actin pedestal, whereas the FHOD1-GFP (green) localized to the base of the pedestals. Actin was stained with Phalloidin-647 (grey). Scale Bar = 5 μm. The inset shows the zoomed region of the pedestals. Scale Bar = 2 μm. **(B)** NIH3T3 cells infected with EPEC for 3 hours (top panel) and 6 hours (bottom panel) are shown. The pedestals showed higher levels of endogenous Arp3 at pedestals 3 hours post infection compared to 6 hours post infection. The arrow points to the site of FHOD1 localized. In contrast, FHOD1-GFP localization to the pedestals was much higher 6 hours post infection compared to 3 hours post infection. The endogenous Arp3 was labeled by mouse anti-Arp3 antibody and detected by anti-mouse AlexaFlour 647 (red). Actin was stained with Phalloidin-568 (grey). Scale Bar = 5 μm. **(C, D)** Quantification of Arp3 and FHOD1 intensity at the pedestal are depicted as box plots. The intensities at the pedestals were normalized to the average fluorescence intensity of the cell. The values are expressed as a.u. (arbitrary units). The graph represents the results of three, independent experiments with 60-80 pedestals used for each group. In the box plots, values of the median (black line), the mean (red dot), lower and upper quartiles (box) and the standard deviation (whiskers) are indicated. The significance of the difference between groups was estimated by two-tailed Student’s *t* test. n.s., P > 0.05 (nonsignificant); **, P ≤ 0.01; ***, P ≤ 0.001; ****, P ≤ 0.0001.

Because the differences in the intensities of Arp3 and FHOD1 were dependent on pedestal size (Fig. S5), we set out to determine whether there were differences in the timing of Arp3 and FHOD1 localization. For example, perhaps Arp3 was active at the initial phases of pedestal formation and FHOD1 was active at a later stage. To test this possibility, we infected NIH3T3 cells with EPEC and fixed them at 3 hours and 6 hours post infection. Arp3 intensity at pedestals was much greater 3 hours post infection (1.75-fold higher) compared to 6 hours, and the trend was inverted for FHOD1 (Fig. 7B-D). Thus, the pedestals were dynamic, heterogeneous structures and their composition changed over time during EPEC infection.

To observe the timing of EPEC pedestal formation, we used live cell imaging to observe a single pedestal as it grew. We imaged a cell which already had small pedestals and followed it over time. FHOD1 intensity increased over time (Fig. 8A and 8C). In contrast, Arp3 intensity decreased over a similar period (Fig. 8B and 8D). The graphs in Fig. 8C-D depict the intensity changes of representative pedestals over time. As observed in Fig. 8C, the pedestal intensity of FHOD1 increased from around 1600 arbitrary units (a.u.) to 2100 a.u. over 45 minutes, while Arp3 intensity decreased from 1600 a.u. to ∼1000 a.u. over two hours (Fig. 8D).

**Fig. 8.**
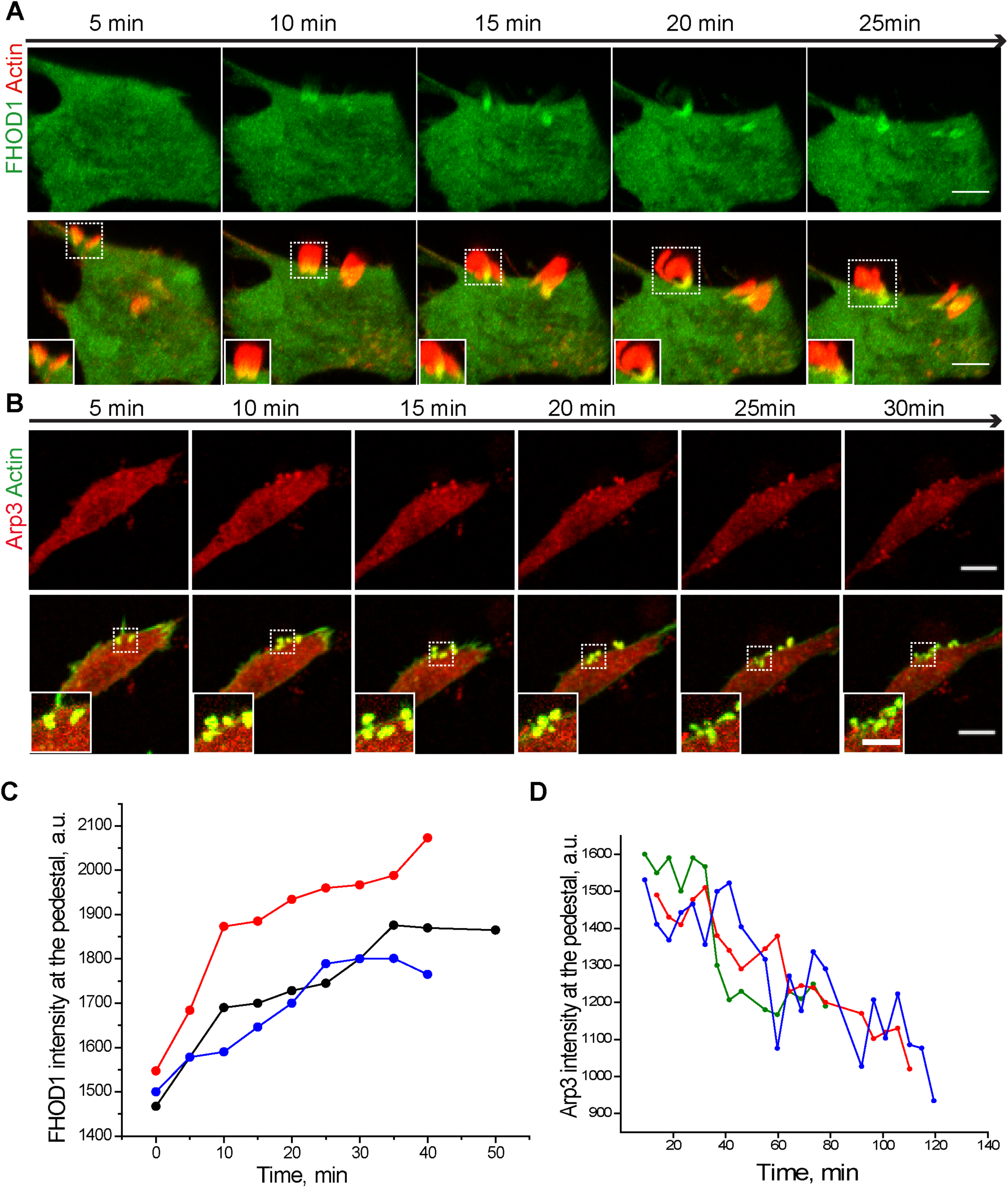
FHOD1 and Arp3 dynamics at an individual pedestal. **(A)** The dynamics of FHOD1-GFP at an EPEC pedestal. The intensity of FHOD1-GFP (green) in the pedestal increased gradually over time. The top panel shows a montage of a time lapse movie of the FHOD1-GFP channel (green), while the bottom panel shows a montage of merged images of the Actin (red) and FHOD1-GFP (green) channel. Scale Bar = 10 μm. A zoomed in region of a pedestal is shown in the inset. Scale Bar = 5 μm. Actin was observed by transfection of Actin-mCherry (red). **(B)** The intensity of Arp-mCherry (red) at the pedestal decreased gradually over time. The top panel shows a montage of a time lapse movie of the Arp3-mCherry channel (red) while the bottom panel shows a montage of merged images of the Actin (green) and Arp3-mCherry (red) channel. Scale Bar = 10 μm. A zoomed in region of a pedestal is shown in the inset. Scale Bar = 5 μm. Actin was observed by transfection with Actin-GFP (green). **(C)** The bleach-corrected intensity (a.u.) of FHOD1-GFP at a single pedestal was plotted over time (three representative pedestals). The intensity of FHOD1 increased almost linearly over time. **(D)** The bleach-corrected intensity (a.u.) of Arp3-mCherry of three representative pedestals over time was plotted. The Arp3 mCherry intensity at the pedestal decreased over time.

## Discussion

### Formin FHOD1 elongates EPEC pedestals

Because formins have been reported to contribute to actin assembly in pathogens (Truong et al., 2013; Alvarez & Agaisse, 2013; Truong, Copeland, & Brumell, 2014; Velle & Campellone, 2018), we explored formin FHOD1 in EPEC pedestals. Using a small molecule inhibitor of formins (SMIFH2), RNAi experiments, and the 3D visualisation and analysis software Imaris, we found that amongst the several formins we tested, formin FHOD1 localised to EPEC pedestals and was involved in pedestal elongation in NIH3T3 cells. That formin FHOD1 affected pedestal size is in agreement with previous finding where siRNA-mediated knockdown of FHOD1 resulted in a smaller volume of lamellipodia-like ruffles caused by *S.* Typhimurium invasion (Truong et al., 2013). Moreover, because the pedestal FHOD1 intensity increased as pedestals elongated, and there were no significant differences in the number of pedestals formed in the absence of FHOD1 (this work; Velle & Campellone, 2018), we propose that FHOD1 is involved in the later stages of pedestal formation.

### Pedestal formation is a two-step process

Pathogens such as *Vaccinia virus, Listeria* and *Shigella* exploit the host cytoskeleton for their intracellular motility facilitated by actin ‘comet tails’. Recent studies have shown that formin FHOD1 promotes the elongation of actin comet tails initiated from Arp2/3 by *Vaccinia virus* (Alvarez & Agaisse, 2013). It is interesting to note that EPEC pedestals also come in different shapes and sizes: some are short, and stub-like, whereas some are very long and curved, and protrusion-like. Based on our results, we propose that EPEC pedestal formation is also a two-step process (similar to *Vaccinia virus*), with the initial phase being Arp2/3-dependent and the elongation phase being formin-dependent.

The multi-step process of EPEC pedestal formation allows for the potential involvement of mechanical force in pedestal motility. Actin nucleators fluctuate temporally at the pedestals, with Arp2/3 present early in pedestal formation, while FHOD1 arrives later (Figs. 7, 8). This fluctuation is likely accompanied by a shift in the three-dimensional actin structure in the pedestals from a purely branched actin network to a parallel actin-dominating network. A modelling analysis proposed that branched or purely parallel actin networks generate different forces. Branched actin networks generate higher forces at the expense of motility, whereas purely parallel actin structures generate little or no force, but can aid in high speed movements (Dmitrieff & Nédélec, 2016). These opposing forces may have consequences during pedestal formation and EPEC pathogenesis. Since EPEC is an extracellular pathogen, strong forces might be required to avoid phagocytosis. Branched actin formed from Arp2/3 complexes with the assistance of formin mDia1 in the pedestals can thus act as a force to oppose phagocytosis at the initial phase of infection (Kage et al., 2017; Velle & Campellone, 2018). At the later stage of infection, small forces might be necessary for EPEC motility. Parallel actin structures formed from formins near the base of pedestals can thus enable and facilitate pedestal motility (up to 0.07 μm/s) (J. M. Sanger, Chang, Ashton, Kaper, & Sanger, 1996). Therefore, a multi-step process with different forces dominating at different times is implied in EPEC pedestal formation.

### EPEC pedestals resemble focal adhesions in Src family kinase (SFK) recruitment of formin FHOD1

FHOD1 is targeted by SFKs to adhesion sites (Koka, Minick, Zhou, Westendorf, & Boehm, 2005; Yu, Law, Suryana, Low, & Sheetz, 2011; Iskratsch et al., 2013). In the present work, we demonstrate that a similar event functions during host-pathogen interactions. In mouse fibroblast cells deficient in Src, Yes and Fyn (SYF^-/-^ cells) (Klinghoffer et al., 1999), the localization of FHOD1 to both the initial focal adhesions and EPEC pedestals was lost, but was restored by the re-introduction of SFKs (Iskratsch et al., 2013; and Fig. 5). The functional redundancy among SFKs was apparent in both focal adhesions and EPEC pedestals (Iskratsch et al., 2013; and Fig. 5). The SFK inhibitor PP2 inhibited actin polymerization from early integrin clusters (Yu et al., 2011), and the enrichment of FHOD1 to clusters was also reduced (Iskratsch et al., 2013). Further, both FHOD1 Y99F and an FHOD1 construct in which the poly-proline region was deleted (Δpoly-Pro) were inactive at clusters with reduced localization of FHOD1 (Iskratsch et al., 2013). As a result, FHOD1 was proposed to interact with SFKs for its targeting to the adhesions and for activation by ROCK (Iskratsch et al., 2013). Similarly, in the case of EPEC pedestals, neither FHOD1 Y99F, nor FHOD1 Δpoly-Pro were localized to actin-rich pedestals (Fig. 4; Shah, 2017). We suggest that FHOD1 Y99 and the poly-proline domains employ a common targeting mechanism to interact with SFKs for correct localizaton to EPEC pedestal sites. These findings highlight the resemblance of EPEC pedestals to focal adhesions in their molecular mechanisms of regulation. Hence, their similarities have been expanded from shared static protein composition such as talin and vinculin, to dynamic molecular interactions (Finlay, Rosenshine, Donnenberg & Kaper, 1992; Goosney, DeVinney & Finlay, 2001; Cantarelli et al., 2001).

In summary, our results identify EPEC pedestal formation as a complex process regulated by temporal and spatial coordination of events involving multiple actin nucleators. Since we observed that formin FHOD1 localization required Tir phosphorylation at Y474, it will now be informative to identify whether an additional EPEC effector drives the pedestal localization of formin FHOD1, and to elucidate its role in activating a formin-dependent pedestal formation pathway in molecular detail.

## Materials and Methods

### EPEC strain

E2348/69 (https://www.uniprot.org/proteomes/UP000008205) and EPEC-GFP (GFP-expressing EPEC; Levine et al., 1978) were used for wildtype EPEC infections. EPEC Y474A, EPEC Y454A and EPEC Y454AY474A (a gift from Gad Frankel; Crepin et al., 2010) were used for EPEC Tir mutants infections.

### Cell culture and infection

NIH3T3 (ATCC CRL-1658) cells were regularly cultured in Dulbecco’s modified Eagle’s medium (DMEM) with 10% fetal bovine serum (FBS) and 1% penicillin/streptomycin at 37°C and 5% CO_2_ to 70%–80% confluency and passaged the day before the experiment. The SYF-/- cells were Src/Yes/Fyn triple KO cells used in (Klinghoffer et al., 1999) and it was a gift from Michael Sheetz lab. They were cultured in conditions similar to the NIH3T3 cells. A single colony of EPEC was inoculated in LB broth overnight at 37°C in reduced shaking conditions. The overnight EPEC suspension was used to infecting NIH3T3 fibroblasts cells. For infection, NIH3T3 Cells were seeded for 24 hours at 37°C and 5% CO_2_ in DMEM with 10% FBS and 1% penicillin/streptomycin. For infection, the overnight grown culture of EPEC was diluted 1:1000 in DMEM (in the absence of antibiotics) and then used for infection such that the final multiplicity of infection (MOI) of 150. The culture dishes were then incubated for 5 hours at 37°C in 5% CO_2_.

### Cell transfection and plasmids

NIH3T3 cells were trypsinized (1X TrypLE Express (Gibco^®^ by Life Technologies), 5-10mins, 37°C) and transfected with GFP-tagged plasmids (FHOD1, FHOD1 Y99F, DAAM1, FMNL2, INF2, mDia1, mDia2) or/and mCherry-tagged plasmids (Arp3, Src, Fyn) using Neon transfection system (In Vitrogen). All of the plasmids used in this study were a gift from the Sheetz lab (Iskratsch et al., 2013).

### Inhibitor treatment and knockdown

The inhibitors CK666 and SMIFH2 were obtained from Sigma. The cells were pre-treated with the indicated concentration of inhibitor for 5 hours in the case of SMIFH2 (25 µM) and for 3 hours in the case of CK666 (50 µM) prior to infection. For post treatment, NIH3T3 cells were infected by EPEC for 5 hours. After a wash with 1XPBS was, DMEM with 100 µM SMIFH2 was added to the cells. This was immediately transferred to the live chamber of the confocal microscope and imaged. The siRNAs were obtained from Dharmacon as a mixture of four individual oligos targeting each gene (Arp3: LQ-046642-01-0002; FHOD1: LQ-057380-01-0002). The siRNAs were transfected using jetPrime transfection reagent (Polyplus Transfection). After incubating for 48 hours, the cells were either infected by EPEC using the protocol as above or analysed for western blot.

### Immuno-staining

Cells were rinsed twice with 1X Phosphate buffered saline (PBS) followed by fixation using 4% paraformaldehyde (Sigma, USA) in 1% PBS Buffer for 10 minutes. The dilutions of 1:1000 and 1: 500 were used for Arp3 and FHOD1 (Santa Cruz Biotechnology) immunostaining. Cells were washed and permeabilized with 0.3% Triton-X (Sigma, USA) in 1X PBS for 10 minutes. After washing twice with 1X PBS, the cells were blocked with 5% BSA (bovine serum albumin) for 1 hour. This was followed by incubation with the required primary antibodies diluted appropriately in the blocking buffer. This was followed by a wash with 1% PBS and incubation with the corresponding secondary antibody (anti-mouse AlexaFlour 647, ThermoFisher SCIENTIFIC) along with Hoechst-33342 (1 mg/ml) or DAPI (ThermoFisher SCIENTIFIC, 1mg/ml) and Phalloidin Alexa Fluor® 488 or 568 (Life Technologies, USA, 1:100) for 1 hour.

### Microscopy

Cells were imaged using Nikon A1R Confocal microscope or Nikon W1 Spinning Disk (super-resolution mode) microscope using the 100X oil immersion objective (1.40 NA, CFI Plan-ApochromatVC) and a Neo sCMOS camera. For Live imaging, NIH3T3 cells were infected 1 hour prior to imaging. The dishes of infected cells were then transferred onto the live chamber of the confocal microscope. The cells were imaged in a complete medium (media with serum) at an acquisition rate from 1 to 5-min intervals for 2 to 3 hours. For both fixed and live imaging, z-depth of 0.5µm was used.

### Image analysis

The surface area and intensity measurement for the pedestals was performed using the software Imaris. The Z stacks of images were reconstructed in 3D in Imaris. The pedestal surfaces were drawn using the ‘surface draw’ tool and the actin channel. The intensities for the corresponding channels were then obtained for the same surfaces. To track the surface area and intensity over time, similar surfaces were drawn at time 0 and the intensities were recorded over time. The number of pedestals per cell was manually counted. The percentage of FHOD1-actins overlapping areas over EPEC pedestal was obtained, firstly, by performing Z projection (SUM, Fiji software) of Z stacks of images and obtaining the areas of EPEC pedestals and FHOD1 along the pedestals by masking the actin channel and the FHOD1-GFP channel separately. Then, AND function (ROI manager, Fiji software) was performed to obtain the overlapping regions of these two. By dividing the overlapping regions between actins and FHOD1, we obtained the percentage plot in Prism7^®^.

### Statistical significance

In the current study, two-tailed Student’s t test was used for comparison between two groups. The surface area is depicted as box plots. Values of median (black line), mean (red dot), lower and upper quartiles (box) and the standard deviation (whiskers) are indicated.

### Immunoblotting

For verification of knockdown experiments, cells were lysed in RIPA lysis buffer (ThermoFischer) 48 h after transfection. Extracted proteins were separated by 4–20% SDS-PAGE (Thermo Fisher Scientific), transferred to PVDF membranes (Bio-Rad Laboratories), incubated at 75 V for 2 h, and blocked for 1 h with 5% non-fat milk (Bio-Rad Laboratories) or BSA (Sigma-Aldrich). The polyvinylidene difluoride (PVDF) membranes were incubated overnight at 4°C with appropriate antibodies: anti-Arp (dilution 1:1,000; Sigma); anti-FHOD1 (dilution 1:1,000; Santa Cruz Biotechnology); After three washes (10 min each), appropriate secondary antibodies conjugated with Horseradish Peroxidase (HRP; Bio-Rad Laboratories) were incubated for 1 h, washed three times (15 min at RT), and detected by Pico Western blotting substrate (Thermo Fisher Scientific).

## Acknowledgements

We are extremely grateful to Gad Frankel, Imperial College London, for the kind gift of EPEC mutant strains. We thank Profs. Alexander Bershadsky, G. V. Shivashankar and Michael Sheetz for the helpful discussions and comments. The authors declare that no conflicts of interest exist.

## Supplementary Figure Legends

**Fig. S1. The effect of SMIFH2 on EPEC growth**

**(A)** The platform used for image analysis is shown. The top panel shows a Z-projected image of an EPEC infected cell. The bottom panel shows the same cell represented using the 3D-reconstruction software Imaris. As shown in the right-most panel, the pedestals formed in all planes can be segmented (yellow). **(B)** A growth curve of EPEC (Optical density at 600 nm) does not show any difference in the presence or absence of high concentrations of SMIFH2. The red line represents 100 µM SMIFH2 and the black line represents growth of EPEC in the absence of inhibitor.

**Fig. S2. The effect of CK666 on EPEC pedestal formation**

**(A)** The top panel shows control cells treated with DMSO only. The left-most panel is EPEC-expressing GFP (green), next to it is Phalloidin 568-stained actin (red) and next to it is shown the merged image. The number of pedestals formed per cell decreased when cells were pretreated with 50 *μ*M CK666. Scale bar = 5 *μ*m. The right most images show the zoomed in image of the lamellipodia (Control) converted to multiple filopodia with CK666 treatment. Scale bar = 2 *μ*m. **(B)** The number of pedestals formed per cell reduced by 50% with CK666 treatment. The mean±S.E.M is indicated. The graph represents results of three independent experiments with in total 80-100 cells used for each group. **(C)** Quantification of the total surface are of the pedestals using Imaris. The total area of the pedestal was unchanged with CK666 treatment compared to the control. The graph represents the results of three, idependent experiments with 100-150 pedestals used for each group. In the box plots, values of media (black line), mean (line dot), lower and upper quartiles (box) and the standard deviation (whiskers) are indicated. The graph represents results of three independent experiments with in total 80-100 cells used for each group.

**Fig. S3. Localization of endogenous FHOD1 to the pedestal**

**(A)** Endogenous FHOD1 localized to the base of pedestals. FHOD1 was labeled with mouse monoclonal antibody detected with Alexa Flour-647 anti-mouse antibody (cyan). Actin (red) was stained with Phalloidin-568 and EPEC-GFP (green) was used for infection. The bottom panel shows a zoomed in image of the EPEC pedestals with FHOD1 localized to them. **(B)** The plot is a line scan across a pedestal. The intensity from the edge of the cell of FHOD1 (cyan) and actin (red) was plotted. The peak of FHOD1 intensity at the pedestal coincides with the peak of actin intensity near the bacteria. **(C)** Endogenous FHOD1 staining was abolished in the FHOD1 siRNA treated cells. The top panel shows strong FHOD1 localization at the pedestal in a control cell treated with non-targeting siRNA. The bottom panel shows the loss of FHOD1 staining at the pedestal in cells treated with FHOD1 siRNA. FHOD1 was labeled with mouse monoclonal antibody detected with Alexa Flour-647 anti-mouse antibody (cyan). Actin (red) was stained with Phalloidin-568 and EPEC-GFP (green) was used for infection.

**Fig. S4. Knockdown of FHOD1 using individual siRNAs**

**(A)** FHOD1 was knocked down using four independent siRNAs, as well as a pool of all the four siRNAs. A Western blot compares FHOD1 levels in the absence (control) and presence of siRNA; GAPDH was used as a loading control. **(B)** Pedestal size was reduced in each individual siRNA knockdown to different extents, depending on the respective amount of FHOD1 expressed. Actin was stained using Phalloidin 568 and GFP-expressing EPEC was used for infection. Scale bar = 5 µm. **(C)** Quantification of the total surface area of the pedestals using Imaris. The surface area was reduced the most with the treatment of siRNA B, C and D. The graph represents the results of three independent experiments with 100-150 pedestals used for each group. In the box plots, the values of the median (black line), the mean (red dot), lower and upper quartiles (box) and the standard deviation (whiskers) are indicated. The significance of the difference between groups was estimated by two-tailed Student’s *t* test. n.s., P > 0.05 (not significant); **, P ≤ 0.01; ***, P ≤ 0.001; ****, P ≤ 0.0001.

**Fig. S5. Arp3 and FHOD1 levels at the pedestals are heterogeneous.**

The two panels illustrate the variation in Arp3 and FHOD1 intensities at the pedestals 5 hours post infection. The top and bottom panel shows pedestals formed on two different cells. The top panel shows a representative cell with high Arp3 (cyan) levels, whereas low FHOD1-GFP (green) levels at the pedestals. The bottom panel shows a representative cell showing pedestals with high FHOD1-GFP (green) intensity, but relatively lower Arp3 (cyan) intensity. The arrow points to the FHOD1 and Arp3 localized. Arp3 was stained using mouse antibody and detected using anti-mouse Alexa Flour-647 antibody. Actin was stained with Phalloidin-568 (Red). Bacteria and host nuclei are stained using Hoechst. Scale bar = 5 µm.

