## Supplementary figures and images for "Formin FHOD1 regulates the size of EPEC pedestals"

### Supplemental Figure 1

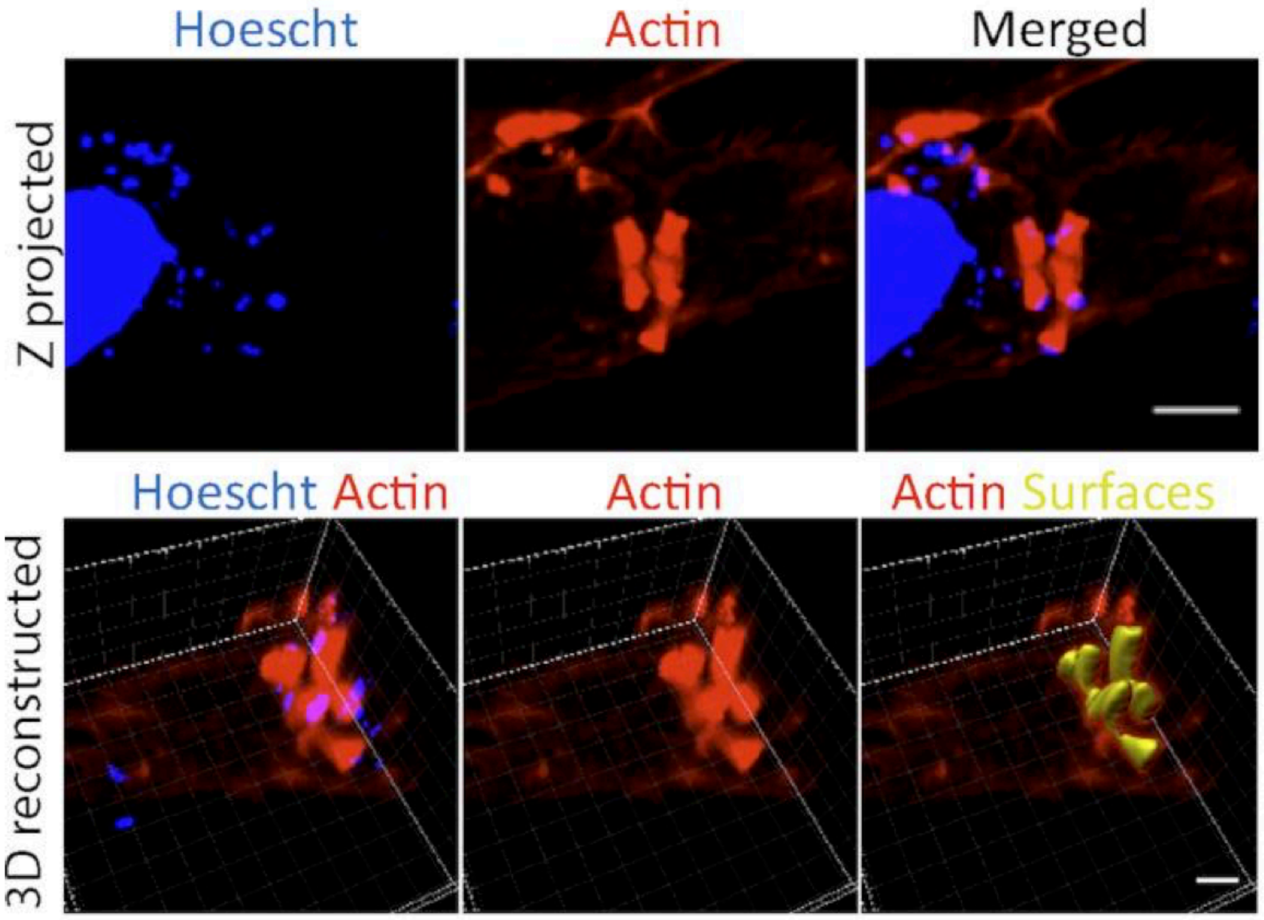

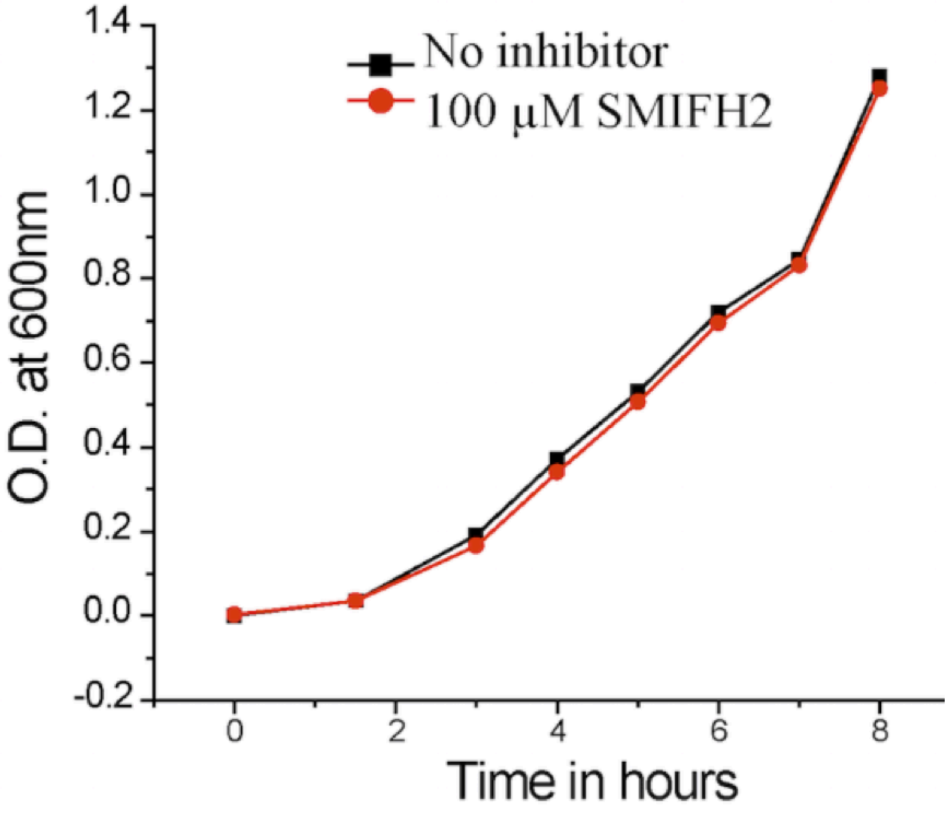

### Supplemental Figure 2

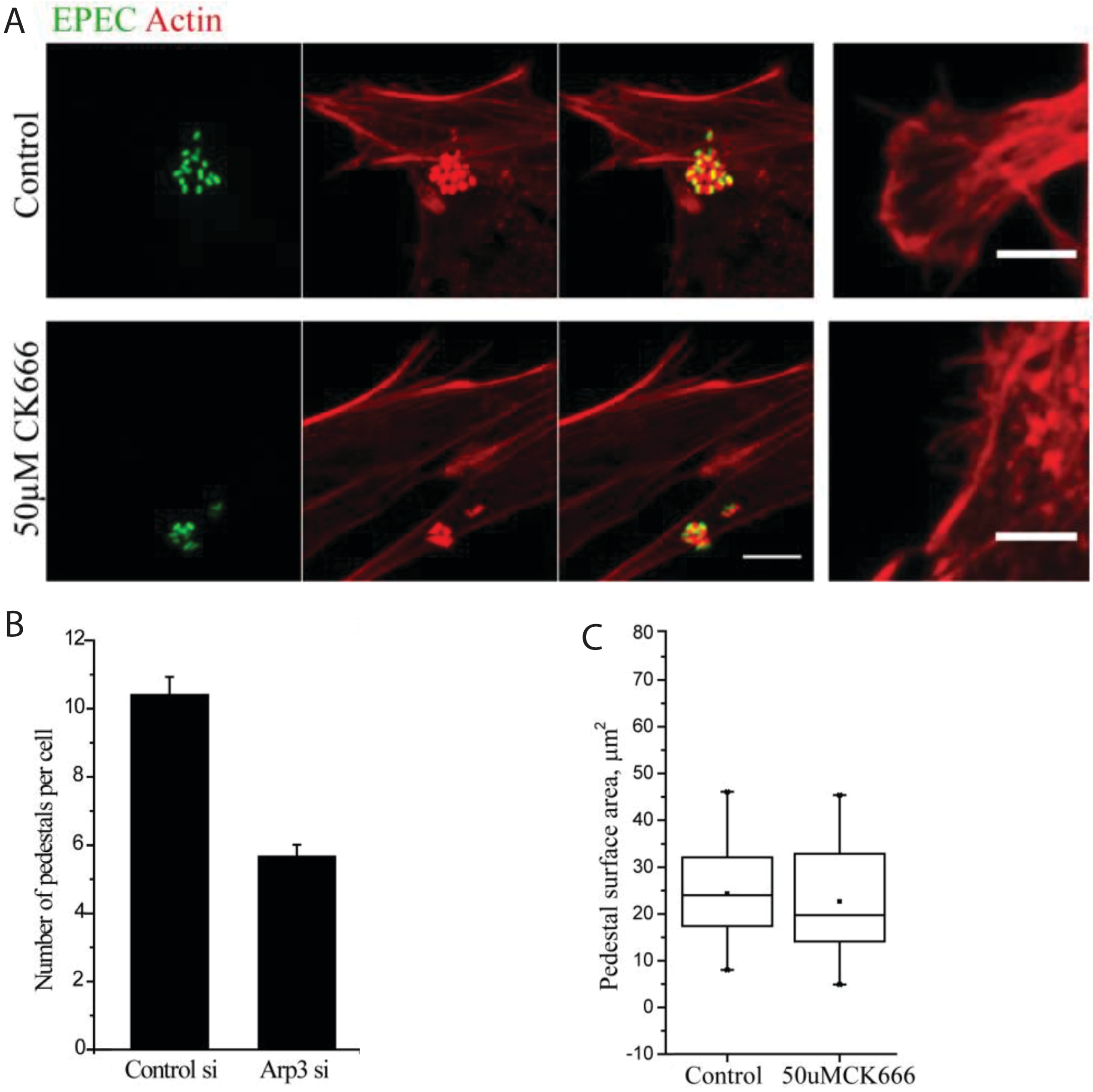

### Supplemental Figure 3

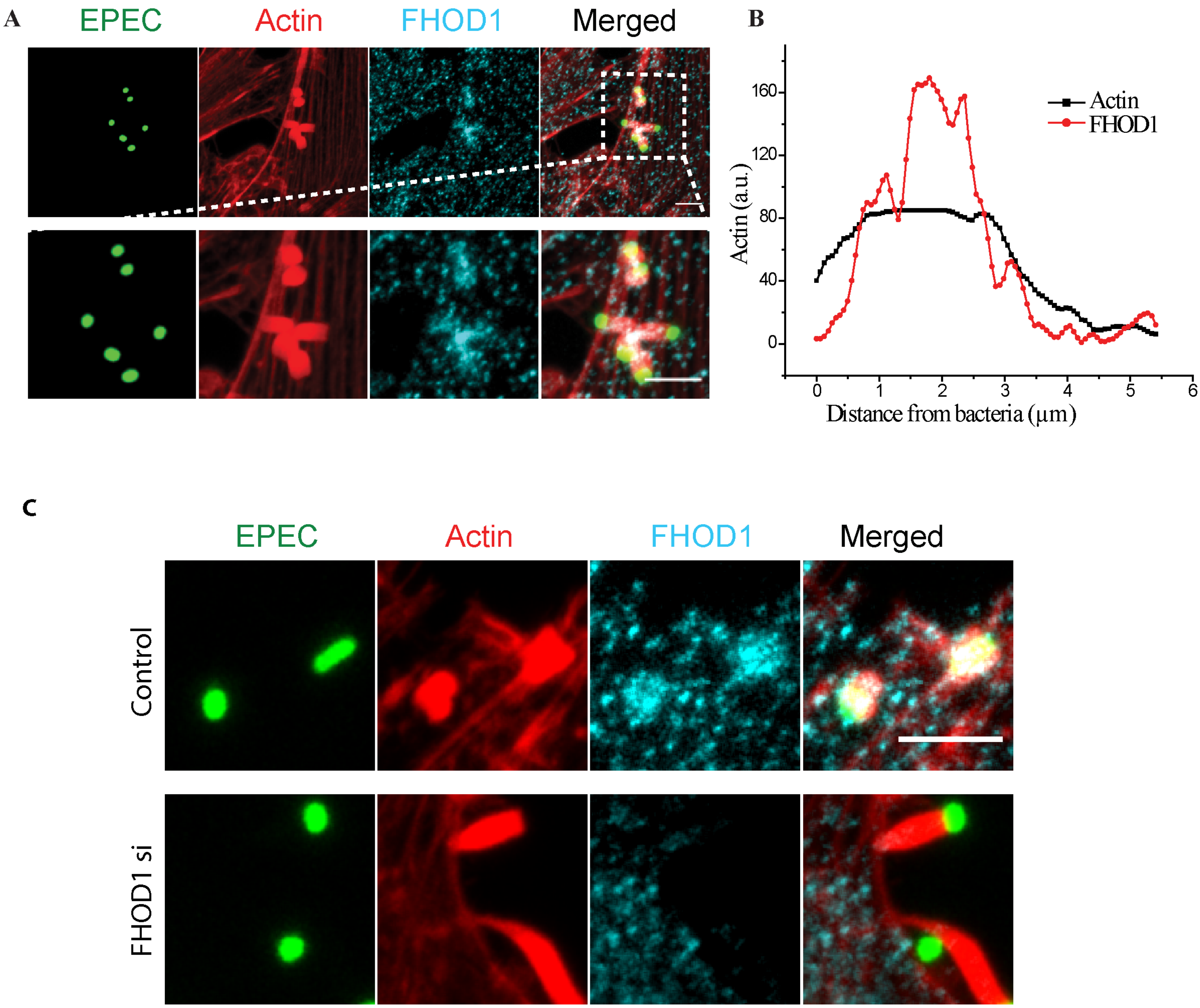

### Supplemental Figure 4

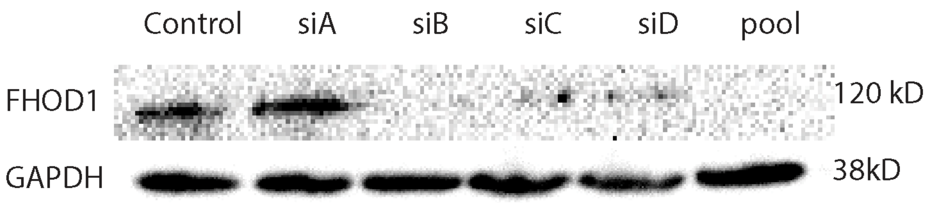

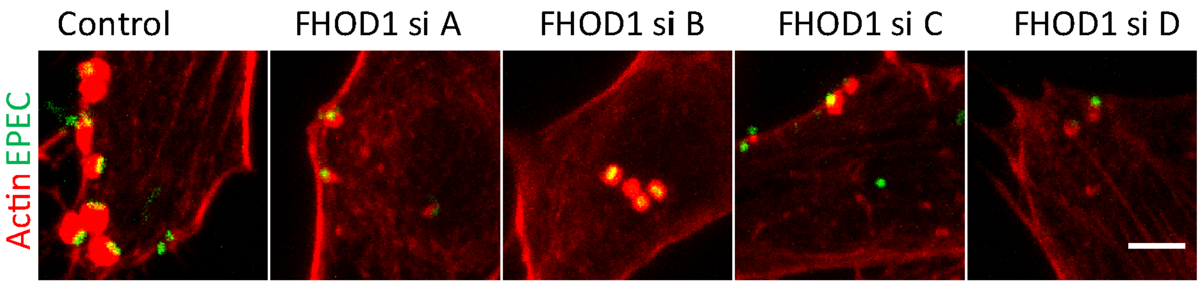

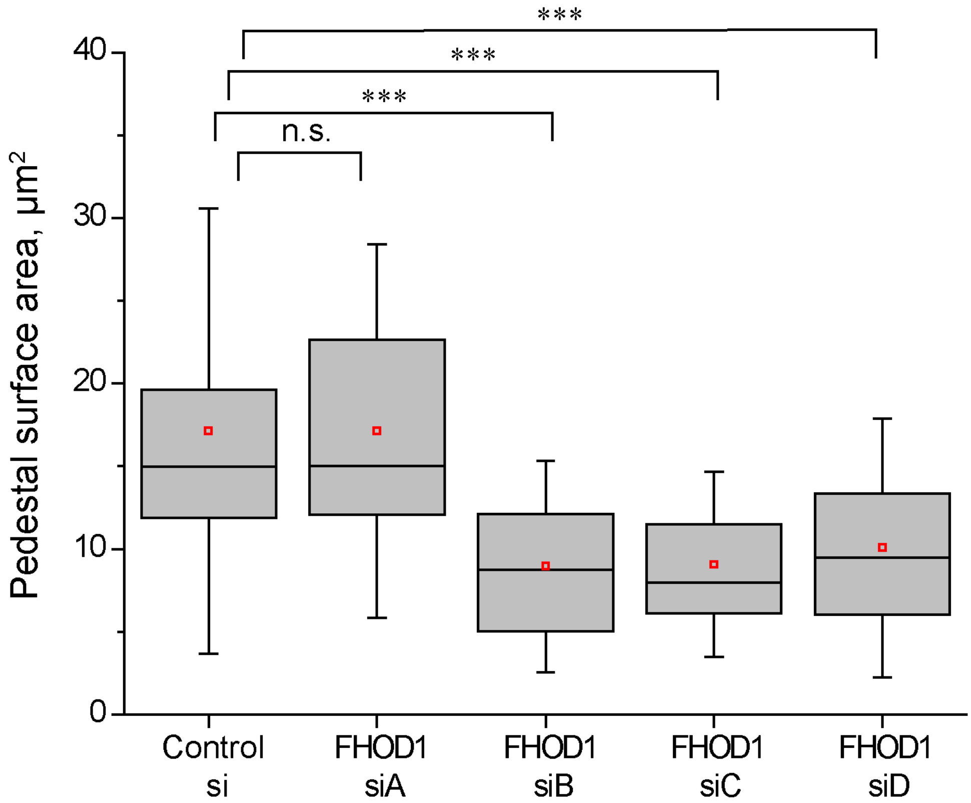

### Supplemental Figure 5

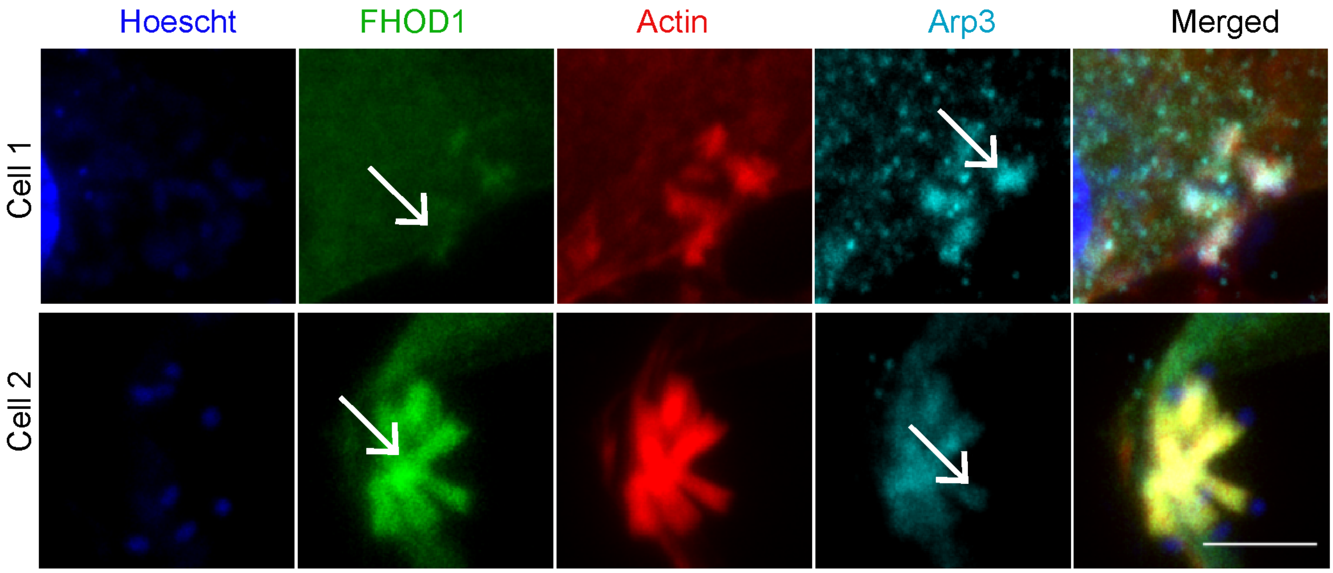
